# Evaluating Tuning Strategies for Sequence Generation with Protein Language Models

**DOI:** 10.1101/2023.02.28.530492

**Authors:** Andrea Nathansen, Kevin Klein, Bernhard Y. Renard, Melania Nowicka, Jakub M. Bartoszewicz

**Author notes:** Equal contribution; corresponding authors. Equal contribution (Andrea Nathansen, Kevin Klein). {, }.

## Abstract

Designing artificial proteins with specialized functions promises new solutions for biological, medical, and environmental use cases. This field benefits from advances in natural language processing, with state-of-the-art text generation models already being successfully applied to protein sequences. Openly available pre-trained protein language models are able to generate artificial protein sequences and can be finetuned on very specific tasks. Considering the high computational cost of finetuning a model exclusively for one downstream task, prompt tuning has been proposed as a more cost-efficient alternative that shares one model across different tasks. However, no openly available implementation of this approach compatible with protein language models has been previously published. Thus, we adapt an open-source codebase designed for NLP models to build a pipeline for prompt tuning on protein sequence data, supporting the protein language models ProtGPT2 and RITA. We benchmark this implementation for generating proteins of a specific family and evaluate the approach using text processing metrics as well as family membership prediction and protein activity prediction of generated sequences. Our results confirm the advantages of prompt tuning in resource usage, especially storage, encouraging further research and expansion of this technique to related use cases. For our evaluated use case, prompt tuning does not reach up to finetuning in terms of the quality of generated protein sequences, indicating the need for more extensive optimization. Lastly, we observe discrepancies between results of similar evaluation tools, highlighting open problems for principled assessment of protein sequence generation quality.

## 1 Introduction

Protein design aims to optimize existing natural proteins for higher efficiency, as well as to generate synthetic proteins with specific functions or properties not found in nature. This opens new avenues for tackling challenging problems, such as advancing cancer immunotherapy (Silva et al., 2019), creating more effective antibodies against SARS-CoV-2 (Shan et al., 2022), and improving the enzymatic degradation of plastic waste (Lu et al., 2022). The function of a protein is determined by its sequence of amino acids folding into a three-dimensional structure, enabling interactions on a biochemical level with surrounding molecules. Inspired by advances in natural language processing (NLP) methods, researchers have successfully developed large language models that operate on protein’s amino acid sequences and can generate entirely new protein sequences. For example, ProtGPT2 (Ferruz et al., 2022) is a decoder-only transformer based on the architecture of the GPT-2 (Radford et al., 2019) large model. Further, Hesslow et al. (2022) developed RITA, a collection of four decoder-only transformer models of different sizes based on GPT-3 (Brown et al., 2020), to examine the impact of model size on performance for protein sequences. The ProGen2 (Nijkamp et al., 2022) models are similar to RITA in terms of model architecture, but their training data distribution differs. Additional examples, like ProGen (Madani et al., 2023) and the enzyme-specific ZymCTRL (Munsamy et al., 2022) model, were trained with a set of conditional control tags. Aside from language models, other work introduces solutions based on diffusion models (Lisanza et al., 2023), equivariant models (Shi et al., 2023), and graph neural networks (Strokach et al., 2020). In this study, we focus on tuning strategies for protein language models, as they explicitly support generation of variable-length sequences.

Training large language models from the ground up requires large datasets and vast computational resources, but pretrained models can be finetuned on specific protein families or properties. Dhodapkar (2023) introduced SpikeGPT2, a finetuned ProtGPT2 model that generates SARS-CoV-2 spike protein sequences to predict potential future mutations. Madani et al. (2023) have also finetuned ProGen on protein families unseen during training. In those cases, one model is trained and stored per task. For large language models with sizes up to billions of parameters, this requires a large amount of storage. However, NLP models such as GPT-3 (Brown et al., 2020) can perform well on specific tasks without model finetuning if given a task-specific prompt in addition to the input. Prompt engineering is a method to manually tune the (natural language) prompt in order to further improve the model’s performance on the task (Brown et al., 2020). Recent work in NLP enhanced this approach by introducing P-tuning (Liu et al., 2021), a method for automatically tuning the embeddings for a given prompt. P-tuning preserves the structure of a human-designed prompt with regard to the positions of trainable, continuous tokens. To create prompts that are more agnostic to the natural language structure of the task description, Lester et al. (2021) proposed learning soft prompts as a set of extra embeddings that are prepended to the embedded input of the model. While still including task-specific tuning, prompt tuning reduces the storage requirements per task from a billion-parameter model to a prompt of ten thousand up to a few million parameters. Lester et al. (2022) have further worked on transferring tuned prompts across models.

Although prompt tuning is a promising direction in NLP, only a few approaches have been proposed to make it available for protein language models and evaluate its potential for improving protein sequence generation. ProGen and ZymCTRL, for example, allow specifying control tags for conditional generation (Madani et al., 2023; Munsamy et al., 2022), but the model has to be pretrained or finetuned to understand a defined set of control tags. In contrast, prompt tuning is a method to tune new task specifications without having to finetune the model. Hesslow et al. (2022) report having tested prompt tuning for RITA, but the authors have not made their code available yet, and provide only a brief description that does not allow complete reproducibility. Further, Wang et al. (2023) proposed a prompt-aware transformer for different tasks such as function prediction and masked language modeling. However, they incorporate prompts in pretraining, so the method cannot be applied to existing unconditional protein language models. He et al. (2022) employed prompt tuning for the chemical-protein interaction classification task, but not for sequence generation. Moreover, Lisanza et al. (2023) proposed the generation of sequence-structure pairs using RoseTTAFold with diffusion which can be conditioned on families and functions, but it requires the implementation of guidance signals. For NLP, Lester et al. (2021) have released an implementation of their prompt tuning approach, but it is not immediately applicable to protein language models like ProtGPT2 or RITA.

In our work, we adapt an open-source implementation of prompt tuning for natural language to learn prompts for conditional protein sequence generation. Our pipeline is compatible with ProtGPT2 and the RITA models. In order to reproduce the experiments conducted by Hesslow et al. (2022), we tune prompts for the RITA models on a held-out protein family and evaluate the performance in comparison to the base model. For a more thorough benchmark, we compare prompt tuning to finetuning for this use case. Aside from perplexity as reported by Hesslow et al. (2022), we further evaluate the quality of generated sequences by computationally predicted protein family membership and measures that were shown by Johnson et al. (2023) to indicate in-vitro activity. Our results show an improvement in perplexity similar to the effect reported in Hesslow et al. (2022), as well as an improvement of positive family membership predictions by ProtCNN (Bileschi et al., 2022). Further, prompt tuning has the advantage of lower computational requirements and orders of magnitude smaller prompt storage size compared to whole models. However, HMMER predictions and activity filters indicate little to no improvement over the untuned model. We observe much better performance for the finetuned model after training for a few epochs only, suggesting finetuning as the preferred tuning method for this use case. However, the potential of mass storage of trained prompts motivates additional optimization, as well as further research for other use cases. Thus, we see our work as a promising starting point for an openly available prompt tuning pipeline for pretrained protein language models, facilitating future studies in this direction. The source code, preprocessing scripts, model checkpoints, and data splits used are available at https://gitlab.com/dacs-hpi/protein-prompt-tuning.

## 2 Methods

### 2.1 Prompt Tuning for sequence generation

Prompt tuning, as introduced by Lester et al. (2021), is an automated approach for tuning task-specific soft prompts that are prepended to a pretrained model’s input at the embedding level. A soft prompt is a continuous matrix ***P*** *∈* R^*m×e*^, with *m* being the number of tokens of the soft prompt and *e* the dimension of the model’s embeddings. Soft prompts can be initialized in various ways, such as a concatenation of randomly sampled embedding vectors of the model’s vocabulary or as a matrix filled with random numbers.

In a sequence generation step *i*, the model computes the probability for the next token ***x***_*i*_ conditioned on the previous tokens *{****x***_1_, ***x***_2_, …, ***x***_*i*−1_*}*. When applying a soft prompt ***P*** at the embedding level, the probability for token ***x***_*i*_ is calculated from *{****P***, ***z***_1_, ***z***_2_, …, ***z***_*i*−1_*}* with ***z***_*j*_ being the embedding vector for token ***x***_*j*_.

For a prompt tuning step (Figure 1), we take a training sequence ***X*** =*{* ***x***_1_, ***x***_2_, …, ***x***_*n*_*}* and predict ***x***_*i*_ for each *i* in [1; *n*] with applying ***P*** . After gradient computation, only ***P*** is updated during backpropagation while the model’s weights remain unchanged. The specific training objective and loss function is determined by the model implementation.

**Figure 1:**
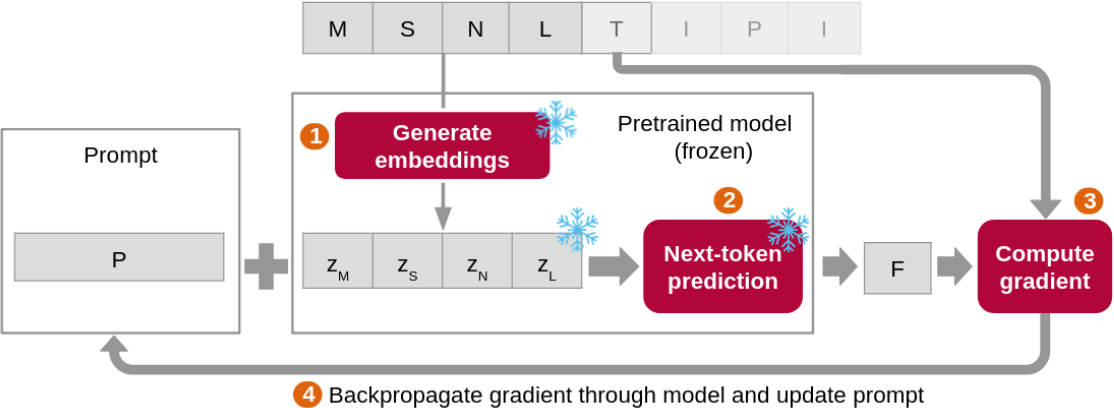
Schematic representation of a prompt tuning step. First (➊), we generate embeddings for the input tokens. Then the prompt is prepended to the embeddings, and the resulting matrix is given as input into the prediction model (➋). Next, a loss function calculates an error for the predicted token w.r.t. the actual next token (➌). This gradient is then backpropagated through the model and updates only the prompt (➍).

### 2.2 Implementation

Lester et al. (2021) published an implementation of prompt tuning^3^ dependent on Jax, Flax, Flaxformer and T5X. However, we employ a codebase relying on Huggingface and PyTorch, as we deem it to be more easily accessible for the wider protein design community. We adapt the independently developed prompt tuning implementation mkultra^4^, built for the NLP language models GPT-2 and GPT-Neo, to build a training pipeline for protein sequences supporting ProtGPT2. Additionally, we add support for the RITA language models to enable a direct comparison with Hesslow et al. (2022). Adaptation to more protein language models like ProGen (Madani et al., 2023) and ProGen2 (Nijkamp et al., 2022) is a subject of future improvement. The pipeline is implemented in Python 3.10 using PyTorch 1.13.1 and Huggingface transformers 4.20.1.

### 2.3 Dataset

Following Hesslow et al. (2022), we conduct our experiments on protein sequences of the Pfam family `PF03272`, which was not included in the pretraining of the RITA models. As the specific dataset used in the experiments of Hesslow et al. (2022) is not available, we construct an analogous dataset consisting of the proteins associated with this family in InterPro^5^ (Paysan-Lafosse et al., 2022). As in Hesslow et al. (2022), we preprocess our data by leaving out sequences containing X (uncharacterized amino acids) and replace the ambiguous characters B, J and Z with a random choice between D and N, I and L, and E and Q, respectively. We create training, validation, and test splits at the ratio of 80 : 10 : 10.

To avoid duplicate sequences between the splits, we cluster the dataset beforehand with MMseqs2 (Steinegger & Söding, 2017, 2018). For each cluster, we select the representative member determined by MMseqs2. Our main dataset is clustered at a 100% sequence similarity threshold to resemble the UniRef100 conditions. It contains 1536 sequences for training, 193 for validation and 193 for testing. We create further datasets by clustering at lower sequence identity thresholds: 95%, (611, 77, and 77 sequences for training, validation, and test, respectively), 65% (283 / 36 / 36), and 35% (191 / 24 / 24). This allows us to evaluate the model’s performance in generating novel protein sequences that are less similar to the sequences seen in training. On the other hand, the datasets become smaller for lower thresholds, which is likely to influence the results as well. More details about the clustering decisions are described in Appendix A.1.

### 2.4 Training Setup

We evaluate our prompt tuning pipeline applied on the RITA models (Hesslow et al., 2022). RITA is an autoregressive decoder-only transformer, trained with the objective of minimizing the cross-entropy loss in next-token prediction. Hesslow et al. (2022) provide four models of different sizes: RITA-S (85M parameters, embedding size 768), RITA-M (300M / 1024), RITA-L (680M / 1536), and RITA-XL (1.2B / 2048) ^6^. We choose RITA-S as the default model size. Similar to the experiments of Hesslow et al. (2022), we run further studies with RITA-M, RITA-L, and RITA-XL to examine the effect of prompt tuning in relation to model size. While we focus on the RITA model family, we also perform a proof-of-principle experiment using ProtGPT2 (see Appendix A.3). Both RITA and ProtGPT2 use the cross-entropy loss function. As described in Section 2.1, in contrast to full model training or finetuning, the model’s weights are frozen and only the prompt weights are updated.

Initializing the prompt weights from the model’s vocabulary embeddings was shown to be superior to random uniform initialization from range [−0.5, 0.5] for NLP models (Lester et al., 2021). We evaluate this in a small ablation study (see Appendix A.4 and Table A1) and conduct our experiments with prompts sampled from the vocabulary due to improved performance for the larger model and almost no change for the smaller one. We set the prompt length to 10 tokens, as suggested by the authors of Hesslow et al. (2022)^7^. Each training runs for a maximum of 300 epochs with early stopping after 20 epochs of no decrease in validation loss. We run each training three times with a different random seed for prompt initialization.

To enable comparison between prompt-tuned and finetuned models, we tune one RITA model for each size with the example finetuning script provided by Huggingface^8^. Following the default settings of the script, we run the finetuning for 3 epochs.

Further training parameters and details about our prompt tuning and finetuning setup are described in Appendix A.2.

### 2.5 Evaluation

To compare sequence generation performance, we measure model perplexity, a common metric for language models equal to the exponential of the model’s cross-entropy loss. Therefore, we expect perplexity to drop for models achieving a better fit to the target task. Hesslow et al. (2022) report perplexity per amino acid to enable comparison between models with different vocabularies. For RITA models, this is equivalent to perplexity per token, as RITA’s vocabulary consists of one token per amino acid. We set a batch size of 2 for this experiment.

Perplexity provides a metric of text generation performance related to the probability of a given sequence, as estimated by the model. However, another level of validation from the biological perspective is necessary. To test whether the produced sequences indeed exhibit the desired biological properties, we predict their family membership with ProtCNN (Bileschi et al., 2022) and measure the percentage of sequences classified as members of the desired family. We set the maximum number of generated amino acids to 1014, which is the model’s context size excluding the prompt tokens (see Appendix A.2). Note that ProtCNN was trained on Pfam seed sequences rather than whole proteins, so we use a sliding window approach to call individual domains in the inputs. Following the procedure described by Bileschi et al. (2022), we use all possible start-end pairs with a minimum length of 50 amino acids as windows for a sequence. Since Pfam family membership is dependent on the presence of specific domains, we count the whole protein sequence as belonging to the target family if ProtCNN predicts the family `PF03272` for at least one window within a sequence. Bileschi et al. (2022) report improved performance for ProtENN, an ensemble of 19 ProtCNN instances. Due to memory and runtime limitations, we conduct a small study for up to 3 instances and a setting where we only count windows with a probability of at least 0.5 (see Appendix A.6). For our evaluations of generated sequences, we use a single model with a probability threshold of 0.5.

Further, we use HMMER v3.3.2^9^ to predict family membership with hmmsearch against the Pfam profile HMM downloaded from InterPro^10^ with the profile-specific gathering thresholds.

To get another indicator for the real-world usefulness of the generated protein sequences, we apply sequence-based filters evaluated in Johnson et al. (2023). These filters were demonstrated to identify active proteins, i.e. proteins that successfully fold in *E. coli* and show their specific catalytic activity in an *in vitro* assay. We evaluate how many generated sequences pass these filters. Appendix A.7 describes the filters in more detail.

In order to compare resource usage between prompt tuning and finetuning, we measure GPU RAM usage, size of the stored models and prompts, and runtime per epoch. Since finetuning runs only 3 epochs, we measure for both prompt tuning and finetuning the average runtime per epoch for 3 epochs, over 2 runs each. For consistent comparison across model sizes, we set a batch size of 1 for all sizes here.

## 3 Results

First, we aim to reproduce the results reported by Hesslow et al. (2022) as the exact setup, code and data used in their experiments are not available at the time of writing. We tune prompts for RITA-S, RITA-M, RITA-L, and RITA-XL and compare the perplexity of each prompt-tuned model to its untuned counterpart on the main test set (clustered at the sequence identity threshold of 100%). In addition, we report performance for the finetuned models, as a comparison between those two tuning strategies has not been done before in the context of protein language models. Similarly to the values presented in Hesslow et al. (2022), prompt tuning leads to a decrease in perplexity (Table 1). This also holds for a smaller experiment using ProtGPT2 (see Appendix A.3). Overall, larger prompt-tuned models perform better. However, perplexity for finetuned models is much lower, with few differences between model sizes M, L, and XL, and L being the best one.

**Table 1:** Perplexity comparison between prompt-tuned model, finetuned and base model (Hesslow et al., 2022) for different model sizes (small, medium, large and extra large), measured on the main test set (clustered at the sequence identity threshold of 100%). For each prompt-tuned model, we report the mean and standard deviation over the three training runs. Both prompt tuning and finetuning improve performance, with larger gain obtained by finetuning.

|  | RITA S | RITA M | RITA L | RITA XL |
| --- | --- | --- | --- | --- |
| **Prompt-tuned model** | $8.73 \pm 0.1$ | $7.25 \pm 0.19$ | $6.01 \pm 0.5$ | $5.96 \pm 0.72$ |
| **Finetuned model** | 1.49 | 1.39 | 1.37 | 1.38 |
| **Base model** | 13.93 | 12.31 | 9.65 | 6.98 |

Next, we tune prompts for datasets clustered with different sequence identity thresholds (Appendix A.1) to evaluate the performance of RITA-S in terms of larger disparities between training and test data (Table 2). As expected, the perplexity of the base model does not fluctuate much between the test sets. The slight differences can be explained by the fact that the clustered datasets contain different samples from the original dataset. The prompt-tuned models show consistent improvement over the base models, although the performance gain decreases noticeably for lower clustering thresholds. This could indicate that learning a prompt successfully generalizing to novel data becomes more challenging with decreased similarity to the training set. However, as the perplexity on the training set follows a similar trend (Table A2), smaller training set sizes for lower thresholds could have influenced the overall fit of the models as well.

**Table 2:** Perplexity comparison between prompt-tuned model and base model for RITA-S on datasets clustered with different sequence identity thresholds (100%, 95%, 65% and 35%), measured on the respective test set. For each prompt-tuned model, we report mean and standard deviation over the three training runs. Performance improvement by prompt tuning decreases for lower sequence identity thresholds.

|  | 100 | 95 | 65 | 35 |
| --- | --- | --- | --- | --- |
| **Prompt-tuned model** | $8.73 \pm 0.1$ | $9.64 \pm 0.02$ | $12.67 \pm 0.08$ | $13.49 \pm 0.07$ |
| **Base model** | 13.93 | 12.99 | 13.83 | 14.24 |

We further evaluate the properties of generated sequences by predicting family membership with ProtCNN (Bileschi et al., 2022) and HMMER and applying protein activity filters (Johnson et al., 2023). For each RITA model size, we generate 193 sequences with a prompt tuned model, a model finetuned on the main dataset, and a base model, respectively. As reported in A.6, our setup of ProtCNN has a precision of 56% and a recall of 99% for sequence classification on the protein family `PF03272`. Compared to the untuned model, prompt tuning improves the percentage of sequences positively classified by ProtCNN (Table 3). Here, the performance decreases for model sizes L and XL compared to sizes S and M, which could indicate that learning prompts for larger models is more difficult. Similar to perplexity, the finetuned model results are still better, with a performance peak for size L and a decrease for size XL. For prompt-tuned models trained on datasets with different clustering thresholds, the percentage fluctuates only slightly (Table 4). The second family prediction method, HMMER, recognizes all sequences from the test set as members of the family, as expected. However, in contrast to the ProtCNN results, none of the sequences generated by the untuned and prompt-tuned models are positively classified by HMMER. For the finetuned models, the percentage ranges from about 25% (size S) to about 55% (size XL). The evaluation of possible protein activity shows similar results: Sequences generated by the prompt-tuned models show little to no improvement over sequences generated by the base models, with less than 10% passing the filters each. Finetuned models, however, perform much better with up to 40% of sequences passing the filters for RITA-XL (Figure 2).

**Table 3:** Percentage of generated sequences classified as belonging to `PF03272` by ProtCNN, compared between prompt-tuned model, finetuned and base model (Hesslow et al., 2022) for different model sizes (small, medium, large and extra large). For each prompt-tuned model, we report mean and standard deviation over the three training runs. Both prompt tuning and finetuning improve performance, with larger gain obtained by finetuning.

|  | RITA S | RITA M | RITA L | RITA XL |
| --- | --- | --- | --- | --- |
| **Prompt-tuned model** | $51.3 \pm 4.08$ | $52.16 \pm 5.51$ | $39.72 \pm 1.36$ | $24.35 \pm 1.53$ |
| **Finetuned model** | 67.36 | 75.13 | 80.31 | 70.47 |
| **Base model** | 9.84 | 11.92 | 13.47 | 16.58 |

**Table 4:** Percentage of generated sequences classified as belonging to `PF03272` by ProtCNN, compared for tuning a prompt on datasets clustered with different sequence identity thresholds (100%, 95%, 65% and 35%). We report the mean and standard deviation each over the three training runs. Base model performance (see RITA-S base model in Table 3) is not influenced by the datasets and therefore not shown. Performance of the prompt-tuned model shows only slight fluctuation for different sequence identity thresholds.

|  | 100 | 95 | 65 | 35 |
| --- | --- | --- | --- | --- |
| **Prompt-tuned model** | 51.3 ± 4.08 | 52.85 ± 3.05 | 53.54 ± 8.94 | 52.16 ± 2.09 |

**Table 5:** Percentage of generated sequences classified as belonging to `PF03272` by HMMER, compared between prompt-tuned model, finetuned and base model (Hesslow et al., 2022). For each prompt-tuned model, we report the mean and standard deviation over the three training runs. Here, only finetuning improves performance.

|  | RITA S | RITA M | RITA L | RITA XL |
| --- | --- | --- | --- | --- |
| **Prompt-tuned model** | 0.0 ± 0.0 | 0.0 ± 0.0 | 0.0 ± 0.0 | 0.0 ± 0.0 |
| **Finetuned model** | 24.87 | 33.16 | 48.7 | 54.92 |
| **Base model** | 0.0 | 0.0 | 0.0 | 0.0 |

**Figure 2:**
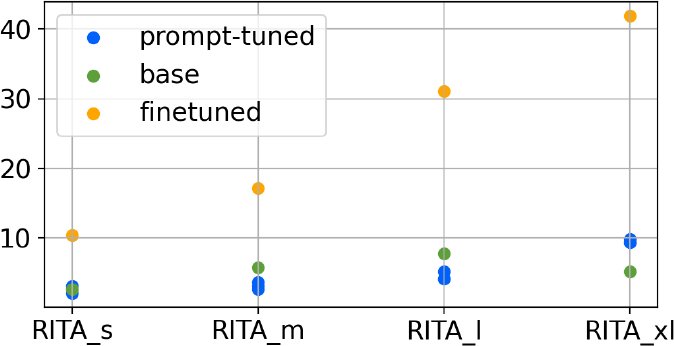
Percentage of sequences passing sequence-based filters for protein activity prediction, compared between prompt-tuned model, finetuned and base model (Hesslow et al., 2022). For each prompt-tuned model, there is one data point per training run.

Additionally, we report computational requirements for prompt tuning and finetuning. The most striking difference is in the required storage size (Table 7). Storing a prompt with a length of 10 tokens requires only 32 to 64 KiB, while the size of the full finetuned model ranges from about 325 MiB (size S) to about 4.5 GiB (size XL). The peak GPU RAM usage (Table 6) is also lower for prompt tuning than for finetuning, with increasing differences for larger model sizes. Further, an epoch for prompt tuning is faster than for finetuning (Table 8). On the other hand, our finetuning ran for only 3 epochs, compared to over 100 epochs for most prompt tuning runs. Thus, finetuning led to better results after much shorter overall training time.

**Table 6:** Peak GPU RAM usage in GiB during the first 3 training epochs, compared between prompt-tuned and finetuned model. Prompt tuning requires less RAM than finetuning, especially for larger models.

|  | RITA S | RITA M | RITA L | RITA XL |
| --- | --- | --- | --- | --- |
| **Prompt-tuned model** | 2.82 | 7.71 | 12.42 | 17.67 |
| **Finetuned model** | 3.70 | 10.60 | 18.44 | 27.95 |

**Table 7:**
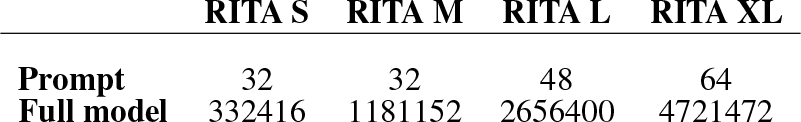
Storage size in KiB for the prompt with a length of 10 tokens compared to the full model. A stored prompt is orders of magnitude smaller in size than a full model.

**Table 8:** Time per epoch in full seconds, measured during the first 3 training epochs for 2 runs, compared between prompt-tuned and finetuned model. A prompt tuning epoch takes less time than a finetuning epoch, especially for larger models.

|  | RITA S | RITA M | RITA L | RITA XL |
| --- | --- | --- | --- | --- |
| **Prompt-tuned model** | $126 \pm 0$ | $310 \pm 0.5$ | $449 \pm 0.5$ | $553 \pm 1$ |
| **Finetuned model** | $127 \pm 0$ | $372 \pm 0.5$ | $675 \pm 0.5$ | $1091 \pm 0$ |

## 4 Discussion

The advantage of prompt tuning lies in its reduced resource usage during training and the greatly reduced storage size of a prompt compared to a full model, especially for large models. With their small size, prompts can be shared a lot easier, e.g. exported and hosted in a small database. There has also been work on recycling prompts between different protein language models (Lester et al., 2022). This motivates the evaluation of prompt tuning for further protein generation use cases, such as the optimization of continuous, rather than categorical properties or combining prompts learned for multiple optimization goals. We share our pipeline with the wider research community for reuse and extension for such further protein generation tasks.

For our evaluated use case, we observe an improvement in perplexity and percentage of positive classification by ProtCNN for the prompt-tuned models. While reproducing the exact setup used by Hesslow et al. (2022) is not possible as the specific design choices, code, and data have not been made publicly available at the time of writing, our results show perplexity improvements consistent with those reported. However, our results show no improvement in HMMER detections and only little improvement in the percentage of sequences passing activity prediction filters. The improvement gained by finetuning, even after much fewer epochs, is much larger. This indicates better suitability of finetuning as a tuning procedure for the task of protein generation for a specific family, at least when training on datasets with high sequence identity threshold. Moreover, contrary to the results shown by Lester et al. (2021), we do not observe a convergence between prompt tuning and finetuning performance for larger models. A possible reason for this could be the difference between natural language and protein semantics, or the fact that Lester et al. (2021) conducted extensive hyperparameter studies while we used default parameters. Large-scale hyperparameter evaluations for prompt tuning on proteins could lead to improved performance.

In our results, we see a noticeable discrepancy between HMMER and ProtCNN. ProtCNN has a low precision in this setup and therefore likely classifies a considerable amount of false positives. On the other hand, given that Bileschi et al. (2022) view their approach as a complement to conventional methods, it is also possible that HMMER has a lower recall on synthetic sequences. Our results highlight the need for employing a variety of metrics for the evaluation of protein generation approaches instead of concentrating on singular measures, such as perplexity only. Future work could include extending our benchmarks, e.g. applying our evaluation setup to more target families and functions, as well as considering structural predictions and experimental validation.

## 5 Conclusion

We present a prompt tuning pipeline for the protein generation models ProtGPT2 (Ferruz et al., 2022) and RITA (Hesslow et al., 2022), adapted from a prompt tuning codebase for NLP. We evaluate the performance of prompt tuning in the use case of protein sequence generation using perplexity, family membership, and activity prediction metrics and compare it to finetuning. Our results show consistently that model finetuning achieves better scores than prompt tuning in in-silico benchmarks while requiring fewer training epochs and hence GPU hours. On the other hand, prompt tuning consumes less RAM and orders of magnitude less storage, making it promising for use cases that were previously economically unviable. Further, while finetuning is already a common technique in protein design, prompt tuning has not been studied in depth for that area. Hence, extensive optimization might close the observed gap in the quality of generated proteins.

Throughout our evaluations, we observe deviations in the results of metrics for generated sequences. These findings underline the necessity of consistent and trusted evaluation metrics for protein language models in general and conditional protein sequence generation in particular.

## Acknowledgments

This work was supported by the de.NBI Cloud within the German Network for Bioinformatics Infrastructure (de.NBI) funded by Bundesministerium für Bildung und Forschung [031A537B, 031A533A, 031A538A, 031A533B, 031A535A, 031A537C, 031A534A, 031A532B] and the Research Program “Designing for Sustainability” funded by the Hasso Plattner Foundation.

## A Appendix

### A.1 Clustering the dataset

The `PF03272` dataset consists of all protein sequences assigned to this family deposited in InterPro. Here, we address the possibility of duplicate sequences in the data, as well as investigate the effects of similarities between the sequences within a single family. We employ MMSeqs2 (Steinegger & Söding, 2017, 2018) to cluster the raw dataset by sequence identity, setting the thresholds between 0% and 100% with a step size of 1%. The number of clusters and the size of the largest cluster for different thresholds are plotted in Figure A1. As expected, the number of clusters drops for a lower threshold, which is particularly visible between thresholds 100% and 90%. Thus, to avoid a radical reduction in dataset size, we employ a threshold of 100% for the data used in the main experiments to remove duplicates while retaining a substantial amount of sequences. We additionally create datasets with thresholds of 35%, 65% and 95% to assess the influence of sequence similarity on the model’s performance. Due to high discrepancies between the cluster sizes, we select the representative member determined by MMseqs2 to form new, balanced datasets. As figure A1 shows, at the threshold of 35%, the size of the largest cluster drops from 489 to 329, while the number of clusters barely increases. A similar drop happens at 95%, although to a smaller extent. Threshold 65% corresponds to the mean of 35% and 95%.

### A.2 Training details

We train the prompts using the Adam optimizer (Kingma & Ba, 2014) with a learning rate of 0.001. Due to limitations in GPU memory, we choose a batch size of 2 in order to maintain the same configuration for model sizes S to L, and a batch size of 1 for RITA-XL. The GPU setup is described in Appendix A.5. The context size is 1024 (as in RITA pretraining (Hesslow et al., 2022)), split into 10 tokens for the prompt and 1014 tokens for the protein sequence. When evaluating the performance of the pretrained model without a prompt, we set the same maximum amount of sequence tokens per context, leading to a context size of 1014 in that case. For generating blocks of the respective size, we split longer sequences and pad shorter blocks with a padding token. We prepend a start token and append an end token to each sequence before splitting. The padding, start and end tokens are specified by the model’s tokenizer.

For finetuning, we set the same batch size as for prompt tuning and keep the default settings of the Huggingface script for the other training hyperparameters. By default, we use the Adam optimizer with a learning rate of 5 *∗* 10^−5^. Due to the absence of prompt tokens, the whole context size of 1024 is used for the protein sequence input.

### A.3 Prompt tuning for ProtGPT2

To showcase the compatibility of our pipeline with ProtGPT2, we conduct a small-scale prompt tuning experiment on the main dataset (clustered at the sequence identity threshold of 100%). Note that the sequences of this dataset might not have been withheld from ProtGPT2 pretraining. However, we deem this setup sufficient for a proof-of-principle. As described in Ferruz et al. (2022), we set a context size of 512 (10 prompt tokens, 502 protein sequence tokens). We use the Adam (Kingma & Ba, 2014) optimizer with a learning rate of 0.0001. The batch size is set to 2, and the prompt size to 10 tokens. The prompt is initialized from the model’s vocabulary embeddings. We run training for a single random seed for 30 epochs with the patience of 20 epochs for early stopping.

Results of this training run show a decrease in perplexity to 1578.53, compared to 4833.37 for the base model. Note that these numbers are not comparable to those reported for the RITA models due to differences in perplexity computation. ProtGPT2 uses a different vocabulary where most tokens consist of more than one amino acid, whereas in RITA, each token corresponds to exactly one amino acid.

**Figure A1:**
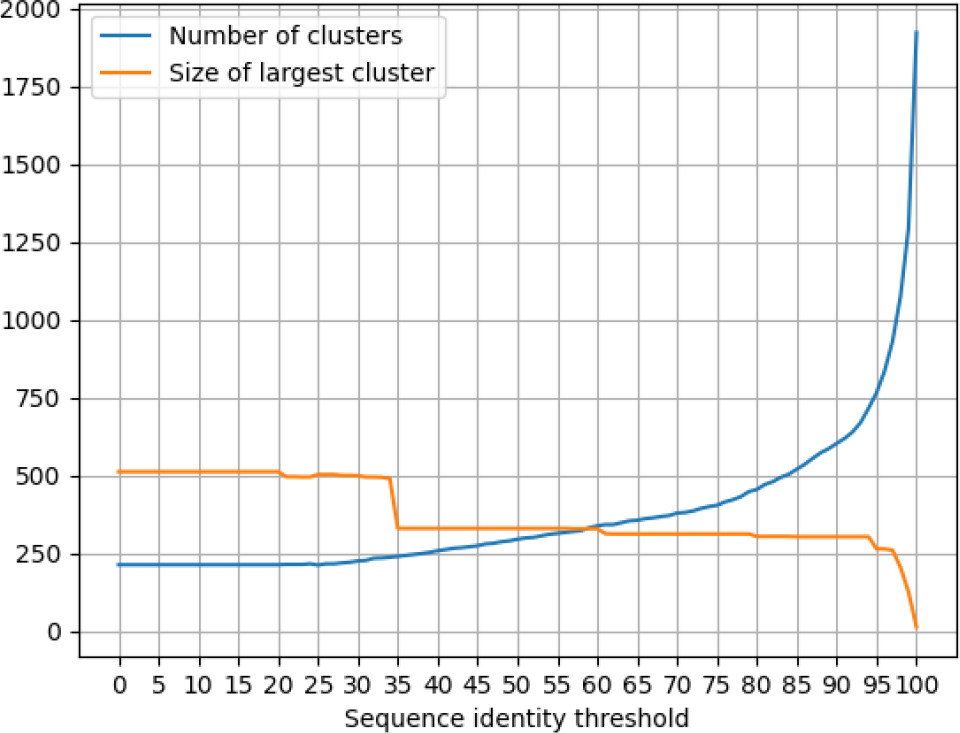
Number of clusters and size of the largest cluster for different sequence similarity thresholds.

**Table A1:** Perplexity comparison between soft prompts with random uniform initialization and soft prompts sampled from the model’s vocabulary for different model sizes (small and medium), measured on the main validation set.

|  | RITA S | RITA M |
| --- | --- | --- |
| **Sampled vocab** | $8.21 \pm 0.08$ | $6.8 \pm 0.17$ |
| **Random uniform** | $8.18 \pm 0.19$ | $7.05 \pm 0.33$ |

### A.4 Ablation study results

To examine whether the findings of Lester et al. (2021) regarding soft prompt initialization hold true for protein language models, we evaluate the performance of soft prompts with random uniform initialization compared to soft prompts sampled from the model’s vocabulary in a small ablation study. We report the perplexity on the main validation set for RITA-S and RITA-M with the different initialization methods in Table A1. For initialization with sampled embeddings, we observe a smaller variance in the results. Using randomly generated vectors leads to minimally better average perplexity for RITA-S, but worse average perplexity for RITA-M. Therefore, we initialize prompts from the vocabulary embeddings in our main experiments.

**Table A2:** Perplexity comparison for the prompt-tuned RITA-S model on datasets clustered with different sequence identity thresholds (100%, 95%, 65% and 35%), measured on the respective training, validation and test sets. For each prompt-tuned model, we report mean and standard deviation over the three training runs. Performance of prompt tuning on all three data splits mostly decreases for lower sequence identity thresholds.

|  | 100 | 95 | 65 | 35 |
| --- | --- | --- | --- | --- |
| **Train** | 8.25 ± 0.08 | 10.13 ± 0.03 | 11.72 ± 0.1 | 11.66 ± 0.12 |
| **Validation** | 8.21 ± 0.08 | 9.65 ± 0.02 | 13.08 ± 0.1 | 12.15 ± 0.06 |
| **Test** | 8.73 ± 0.1 | 9.64 ± 0.02 | 12.67 ± 0.08 | 13.49 ± 0.07 |

**Table A3:** Percentage of sequences classified as belonging to `PF03272` with different ProtCNN configurations, reported for the test set (positive control dataset) and a shuffled version of the test set as well as sequences from other protein families (negative control datasets). The configuration of a single model with the probability threshold is the most accurate one.

|  | Test Set | Other Families | Shuffled |
| --- | --- | --- | --- |
| **Single** | 99.0 | 24.2 | 89.1 |
| **Ensemble 2** | 98.4 | 18.9 | 83.4 |
| **Ensemble 3** | 98.4 | 21.6 | 82.9 |
| **Single thresholded** | 99.0 | 13.2 | 65.3 |

### A.5 GPU setup

Each training and evaluation run was conducted on one GPU at a time. We used a set of NVIDIA A100 GPUs with 40 and 80GB memory. Training took about 10h, 30h and 40h per run for RITA-S, RITA-M and RITA-L, respectively.

### A.6 ProtCNN study

To determine whether ProtCNN performance for the family `PF03272` improves with increasing number of ProtCNN instances, we evaluate consensus predictions (i.e. predictions where all model instances agree on the predicted family for a given window) of ensembles of 2 and 3 instances on the test set. To test for false positives in this method, we apply it on two negative control datasets as well. The first one is derived from the test set with each protein sequence shuffled while preserving monoresidue composition, using esl-shuffle from the HMMER toolkit. The second one is composed of random samples of 10 proteins from each of the 19 other protein families held-out during RITA pretraining^11^. We compare the ensemble performances to the performance of a single model and a setting with a single model applying a probability threshold of 0.5.

As shown in table A3 and table A4, ensemble consensus predictions reduce the amount of predicted false positives as well as predicted true positives. Using a single model with probability threshold of 0.5 leads to a larger increase in precision without a decrease in recall. Therefore, we perform all evaluations of generated sequences with this configuration.

### A.7 Sequence-based filters for protein activity indication

Johnson et al. (2023) report a framework for enzyme activity prediction. They evaluate several computational metrics based on sequence properties, MSA, and structure, on their ability to predict activity. We adopt the sequence-based metrics to gain indication about possible activity of the protein sequences generated by our pipelines. However, we do not include the higher-level evaluation metrics Johnson et al. (2023) integrated into their final framework due to the long runtime of 1-2 hours per sequence on an NVIDIA A100 GPU.

**Table A4:** Precision and recall for the protein family `PF03272` with different ProtCNN configurations, computed from a positive control dataset containing the results on the test set (193 sequences) and a negative control datasets containing a shuffled version of the test set as well as sequences from other protein families (193 + 190 sequences). The configuration of a single model with the probability threshold is the most accurate one.

|  | Precision | Recall |
| --- | --- | --- |
| **Single** | 0.47 | 0.99 |
| **Ensemble 2** | 0.49 | 0.98 |
| **Ensemble 3** | 0.49 | 0.98 |
| **Single thresholded** | 0.56 | 0.99 |

The sequence-based filters include three basic quality checks to rule out unrealistic sequences: The sequences must start with methionine, must not contain long repeats (not more than 3 consecutive identical amino acids or 4 similar tuples) and must not have (predicted) transmembrane domains. For the latter we use *Phobius*, an online tool introduced by Käll et al. (2007) and hosted by Stockholm Bioinformatics Centre of Stockholm University^12^.

Lastly, the sequences are evaluated using the log-likelihood scores emitted by ESM-1v (Meier et al., 2021). Only sequences that score better than 90% of our main test set pass this filter. ESM-1v cannot handle sequences longer than 1024 amino acids and sequences containing ambiguous amino acids, therefore we count those as failed.

## Footnotes

3 https://github.com/google-research/prompt-tuning

4 https://github.com/corolla-johnson/mkultra/commit/a25c72d47980a767b6178861a436900fd83c058f

5 https://www.ebi.ac.uk/interpro/entry/pfam/PF03272/protein/UniProt/, downloaded on January 5, 2023

6 https://huggingface.co/lightonai/RITA_s, https://huggingface.co/lightonai/RITA_m, https://huggingface.co/lightonai/RITA_l, https://huggingface.co/lightonai/RITA_xl

7 https://github.com/lightonai/RITA/issues/3#issuecomment-1131381833

8 https://github.com/huggingface/transformers/tree/v4.20.1/examples/pytorch/language-modeling

9 http://hmmer.org/

10 https://www.ebi.ac.uk/interpro/entry/pfam/PF03272/curation/

11 PF03492, PF03938, PF04153, PF04420, PF04680, PF05318, PF06173, PF06905, PF06917, PF10696, PF10841, PF11968, PF12325, PF12378, PF15340, PF17055, PF17724, PF17988 and PF18369; downloaded on June 2, 2023 via https://www.ebi.ac.uk/interpro/entry/pfam/<family>/protein/UniProt/, where <family> corresponds to the aforementioned family ids.

12 https://phobius.sbc.su.se/

## Notes

### Competing Interest Statement

The authors have declared no competing interest.

### Summary of Updates

Extended evaluation

